# Climate-driven vegetation characteristics shape phytophagous and carnivorous insect diversity in South African savannahs

**DOI:** 10.1101/2024.07.16.603689

**Authors:** Fernando P. Gaona, Sylvain Delabye, Pavel Potocký, Valeriy Govorov, Jan Čuda, Llewellyn C. Foxcroft, Rafał Garlacz, Martin Hejda, Sandra MacFadyen, Tomasz Pyrcz, Klára Pyšková, Ondřej Sedláček, David Storch, Petr Pyšek, Robert Tropek

## Abstract

Despite the recognized importance of insects in savannah ecosystems, the drivers of their diversity patterns remain poorly understood, particularly in the Afrotropical region. This study addresses this gap by investigating the impacts of climate, habitat, disturbance, and vegetation variables on the diversity of phytophagous moths (Lepidoptera) and carnivorous mantises (Mantodea) across 60 plots in Kruger National Park, South Africa. Based on an extensive dataset of 65,593 moth individuals representing 817 morphospecies and 3,511 mantis individuals representing 38 morphospecies, our results revealed plant communities as the fundamental driver of diversity for both insect groups. The effects of vegetation on insect diversity were indirectly influenced by climate, particularly mean temperature (negatively correlated with precipitation), through its impact on plant species richness. Additionally, a complex interplay among bedrock type, water availability, and disturbances from large herbivores and fire further shaped insect diversity. Our findings highlight the region’s vulnerability to climate change, as decreasing precipitation and increasing temperatures were shown to alter vegetation composition and biomass, consequently affecting insect communities. Overall, our results emphasize the necessity of managing large herbivores and regulating fire regimes to maintain diverse vegetation, which is crucial for supporting insect diversity. Effective conservation strategies should prioritize balancing water availability and disturbance intensity, particularly in maintaining the health of seasonal rivers, to mitigate the adverse effects of climate change on these ecosystems.

## Introduction

Biodiversity patterns are shaped by numerous environmental factors, including climate, resource availability, and biotic interactions (Gaston et al., 2000; Wisz et al., 2013; Peters et al., 2016). In savannah ecosystems, large herbivores, fire disturbances, and climatic conditions belong among the fundamental biodiversity drivers across spatial and temporal scales (Pickett et al., 2003; Boogs & Inouye, 2012; Gaget et al., 2020; Araújo & Oliveira, 2021). Previous research on savannah biodiversity has predominantly focused on plants (e.g., Staver et al., 2017; Hejda et al., 2022) and vertebrates (e.g., Ogada et al., 2008; Davies et al., 2016), often overlooking the drivers of insect diversity, despite their crucial roles as pollinators, prey, predators, herbivores, and decomposers (e.g., Dennis et al., 2008; Davies et al., 2012; Castagneyrol et al., 2017; Tonelli et al., 2018).

In savannah ecosystems, plant communities play a key role in determining insect distribution and diversity (e.g., Leal et al., 2016; Parker et al., 2023). According to the resource specialization hypothesis, high plant diversity provides resources for more herbivorous insect species (e.g., Du et al., 2020). Additionally, diverse plant communities create a range of microclimatic conditions and niches that benefit both herbivorous and non-herbivorous insect groups (e.g., Price, 2002; Randlkofer et al., 2010). Numerous studies have demonstrated a positive relationship between plant diversity and various phytophagous insect taxa, including butterflies (e.g., Leone et al., 2023), grasshoppers (e.g., Gebeyehu & Samways, 2002), and omnivorous insects such as ants (e.g., Mauda et al., 2018). Similarly, this relationship extends to predatory and parasitic arthropods (e.g., Procheş & Cowling, 2006; Botha et al., 2017). However, some studies have been less conclusive (e.g., Koricheva et al., 2000; Scherber et al., 2006; Axmacher et al., 2009; Prather et al., 2020; Smith & Willians, 2023), underscoring the importance of abiotic factors, beyond just plant diversity, in explaining of insect diversity patterns (Axmacher et al., 2009; Smith & Willians, 2023).

Thus, the interplay between insect and plant diversity in savannahs is intricate, being influenced directly or indirectly by climatic conditions, soil chemistry, water and nutrient availability, or disturbances from large herbivores and fire (e.g. Shi et al., 2018; Li et al., 2021). As small-bodied ectotherms, insects are particularly sensitive to environmental temperature which is assumed to drive their species richness patterns through physiological constraints in harsh climate (Currie et al., 2004; Kaspari et al., 2019) or temperature-dependent resource exploitation (Brown, 2014). While some previous studies have identified temperature as a primary driver of savannah insect diversity at global (e.g., Dunn et al., 2009), but also at regional scales (e.g., Corcos et al., 2018; Chesters et al., 2019), others have found temperature to be an unreliable predictor, noting instead that rainfall strongly correlates with insect species richness (Parr et al., 2004; Vasconcelos et al., 2018). For example, Delabye et al. (2022a) described a significant increase in moth diversity along a longitudinal gradient of environmental productivity in South African savannahs, predominantly influenced by precipitation but rather independent on temperature in this region. Zhu et al. (2014) experimentally revealed a decline in species richness and abundance of many insect groups in response to both increased and decreased precipitation in a meadow steppe in China. Conversely, Andersen et al. (2015) reviewed that ant diversity in tropical savannahs correlates globally with both temperature and rainfall.

Disturbances from fire and large vertebrate herbivores also drive biodiversity patterns in savannahs (Joern & Laws, 2013). These disturbances not only lead to direct mortality of insects during fires but also indirectly affect its diversity across trophic levels by altering plant growth, community structure, and diversity (e.g. van Klink et al., 2015; Osborne et al., 2018; Mukwevho et al., 2023, 2024). Research on the impact of large herbivores and fire on insect diversity has yielded varied results, reflecting the different responses among insect groups to these disturbances (e.g., Mukwevho et al., 2023). For example, beetles may respond negatively (e.g., Jonsson et al., 2010; Reinhard et al., 2019) while grasshoppers and butterflies might exhibit positive responses to the habitat opening (Joubert et al., 2016; Bonnington et al., 2008; Pryke et al., 2016; Topp et al., 2022). The degree of these responses can vary by influenced by the intensity, frequency, and yearly variation of the disturbances (Jonsson et al., 2010; Coetsee et al., 2019; Korell et al., 2021; Mukwevho et al., 2023, 2024).

This study is a part of the Monitoring Savannah Biodiversity in the Kruger National Park project (MOSAIK), aimed at identifying the principal environmental variables driving the spatiotemporal dynamics of savannah ecosystems and their impact on biodiversity across trophic levels (e.g. Pyšek et al., 2020, Hejda et al., 2022, Čuda et al., 2024). It specifically examines the diversity of moths (Lepidoptera) representing phytophagous insects, and mantises (Mantodea) representing carnivorous insects. Using Structural Equation Models (SEM), we investigate both the direct and indirect effects of climatic and environmental variables on insect species richness (see Figure 1a for the conceptual framework) and their influence on insect community composition.

**Figure 1.**
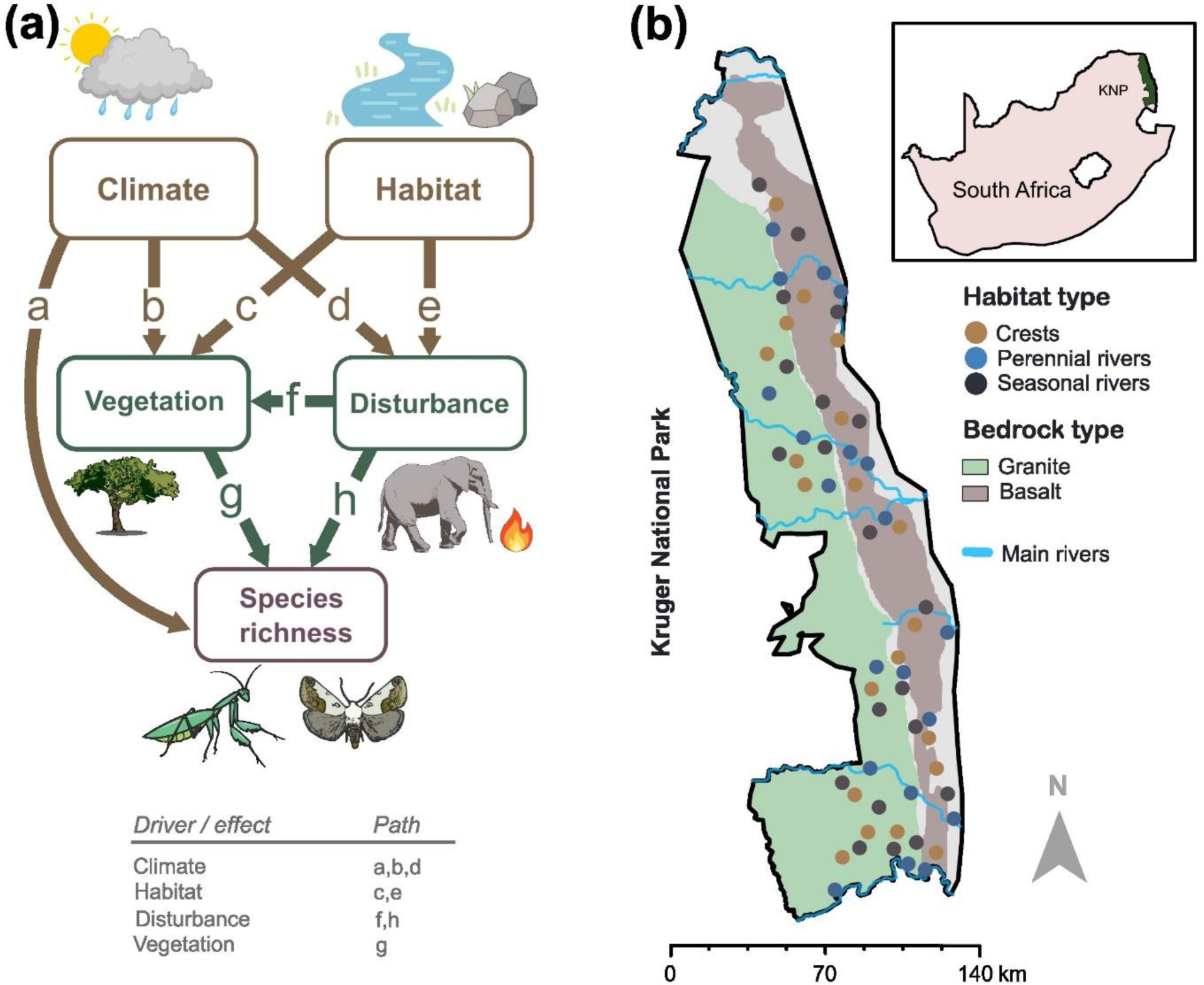
(a) Conceptual framework illustrating the potential direct and/or indirect influence of particular groups of variables on insect species richness. Arrows depict pathways and letters denote different paths aligned with the hypotheses outlined in Introduction. Climate variables include temperature and rainfall; abiotic habitat characteristics comprise water availability and bedrock type; vegetation is characterized by plant species richness, community composition, and biomass; disturbance is represented by the abundance of large herbivores and fire history. All variables are detailed in Table 1. (b) Map of Kruger National Park, South Africa, indicating the 60 study plot locations.

**Table 1.** Environmental characteristics of the studied plots in Kruger National Park. Variables in bold were used in the analyses after the forward selection. Data were taken from Funk et al. (2015)^1^, Wan et al. (2015)^2^, Chuvieco et al. (2018)^3^, Didan (2021)^4^ and Hejda et al. (2022)^5^. Climate and fire-related disturbance variables refer to values per 4 km^2^ grid cell recorded over 20 years (2000–2019). EVI variables refer to values recorded for 250 x 250 m grid cells recorded over the same period. All other variables refer to the 50 × 50 m studied plots and were sampled specifically for the MOSAIC project.

|  | Variable | Description |
| --- | --- | --- |
| Climate | Rainfall (mean) <sup>1</sup> | Average rainfall (mm). |
|  | Rainfall (SD) <sup>1</sup> | Standard deviation of rainfall (mm). |
|  | **Temperature (mean) <sup>2</sup>** | Average environmental temperature (°C). |
|  | Temperature (SD) <sup>2</sup> | Standard deviation of environmental temperature (°C). |
|  | Temperature (max) <sup>2</sup> | Average minimum temperature (°C). |
|  | Temperature (min) <sup>2</sup> | Average maximum temperature (°C). |
| Habitat | **Water availability** | Seasonal availability of water: perennial river, seasonal river, crest. |
|  | **Bedrock type** | Basalt, granite. |
| Disturbance | **Number of fires (mean) <sup>3</sup>** | Average number of fires. |
|  | Number of fires (SD) <sup>3</sup> | Standard deviation of the number of fires. |
|  | **Large herbivore abundance** | Abundance of large mammal herbivores. |
| Vegetation | **EVI (mean)** <sup>4</sup> | Average Enhanced Vegetation Index (EVI). |
|  | EVI (SD) <sup>4</sup> | Standard deviation of EVI. |
|  | **Vegetation composition (PC1)** <sup>5</sup> | 1<sup>st</sup> principal component axis of plant community composition. |
|  | **Vegetation composition (PC2)** <sup>5</sup> | 2<sup>nd</sup> principal component axis of plant community composition. |
|  | Vegetation composition (PC3) <sup>5</sup> | 3<sup>rd</sup> principal component axis of plant community composition. |
|  | Vegetation composition (PC4) <sup>5</sup> | 4<sup>th</sup> principal component axis of plant community composition. |
|  | **Plant species richness** <sup>5</sup> | Total number of plant species. |
|  | Grass species richness <sup>5</sup> | Number of grass species. |
|  | Herb species richness <sup>5</sup> | Number of species in the herbaceous layer (%). |
|  | Shrub species richness <sup>5</sup> | Number of species in the shrub layer (%). |
|  | Tree species richness <sup>5</sup> | Number of species in tree layer. |
|  | Plant cover <sup>5</sup> | Total plant cover (%). |
|  | Tree cover <sup>5</sup> | Tree layer cover (%). |
|  | Shrub cover <sup>5</sup> | Shrub layer cover (%). |
|  | Herb cover <sup>5</sup> | Herbaceous layer cover (%). |

We hypothesise that (1) climatic drivers affect insect species richness both directly (Figure 1a, path a) and indirectly through changes in vegetation (Figure 1a, path b). Specifically, we predict that insect species richness decreases with higher temperatures, surpassing the thermal maxima of some species, and increases with higher precipitation which enhances plant diversity in Kruger National Park (KNP) (Hejda et al., 2022). We also expect that (2) the abiotic habitat characteristics (water availability and bedrock type) indirectly affect species richness of both insect groups through their influence on vegetation (Figure 1a, path c) and disturbance drivers (Figure 1a, path e). We expect that moderate water availability will support the highest insect species richness, due to a sparser presence of large herbivores (Young et al., 2009) which supports higher plant diversity (Hejda et al., 2022). We also expect higher insect diversity on basaltic bedrock, owing to its known higher soil fertility supporting plant diversity, large herbivore abundance, and fire frequency compared to granite soils (Mills & Frey, 2005; Smit et al., 2013; van Wilgen et al., 2022; Hejda et al., 2022). Given the lack of a clear mechanism by which water availability or bedrock type could directly influence insect diversity in KNP, we did not include this direct relationship into the SEM analyses. Furthermore, we expect that (3) fire and large herbivore disturbances mainly have an indirect effect on insect species richness through their impacts on vegetation structure and composition (Figure 1a, path f). We predict a unimodal response of insect species richness, peaking in sites with a moderate level of disturbances. Finally, we expect (4) strong direct effects of vegetation characteristics on both insect groups’ species richness (Figure 1a, path g), which will be strongly correlated to plant species richness and biomass. Given the closer relationship of phytophagous insects to their host plants, we predict a stronger correlation of plant diversity with the diversity of moths than mantises. Similarly, we expect that (5) the community composition of phytophagous and carnivorous insects will be shaped by abiotic habitat characteristics, climate, and vegetation variables. We predict that vegetation variables will be more important predictors of insect community composition than climatic or disturbance variables for both insect groups, although we expect this to be more pronounced for phytophagous moth communities.

## Materials and Methods

### Study area and plots

The study was carried out in Kruger National Park (KNP) (Figure 1b), one of South Africa’s largest and oldest national parks, which hosts diverse flora and fauna. Spanning approximately 20,000 km^2^, KNP features a gradient of subtropical and tropical climates. It is traversed by five major rivers, Sabie, Olifants, Crocodile, Letaba, and Luvuvhu (Figure 1b). Daily temperature ranges from 5.6°C in winter to 32.6°C in summer, and the annual precipitation ranges from 450 mm in the northern parts to 750 mm in the southern parts of KNP (Siebert & Eckhardt, 2008; Chadwick et al., 2013; Appendix S1: Figure S1). The park’s vegetation includes various savannah types, with a variety of dominant woody species and varying proportions of woodlands and open grasslands (du Toit et al., 2003).

Within the MOSAIK project, 60 plots (50 × 50 m) were strategically established across KNP (Figure 2b). They were spatially organized into triplets, with individual plots representing three different levels of water availability: (i) perennial rivers or other permanent water sources, (ii) seasonal rivers which usually dry out during the dry seasons, and (iii) dry crests located at least 5 km from any water source (see Pyšek et al., 2020; Hejda et al., 2022 for more details). Each triplet was situated on either granitic (Skukuza and Phalaborwa landsystems) or basaltic bedrock (Satara and Letaba landsystems) (Figure 1b; Pyšek et al., 2020).

**Figure 2.**
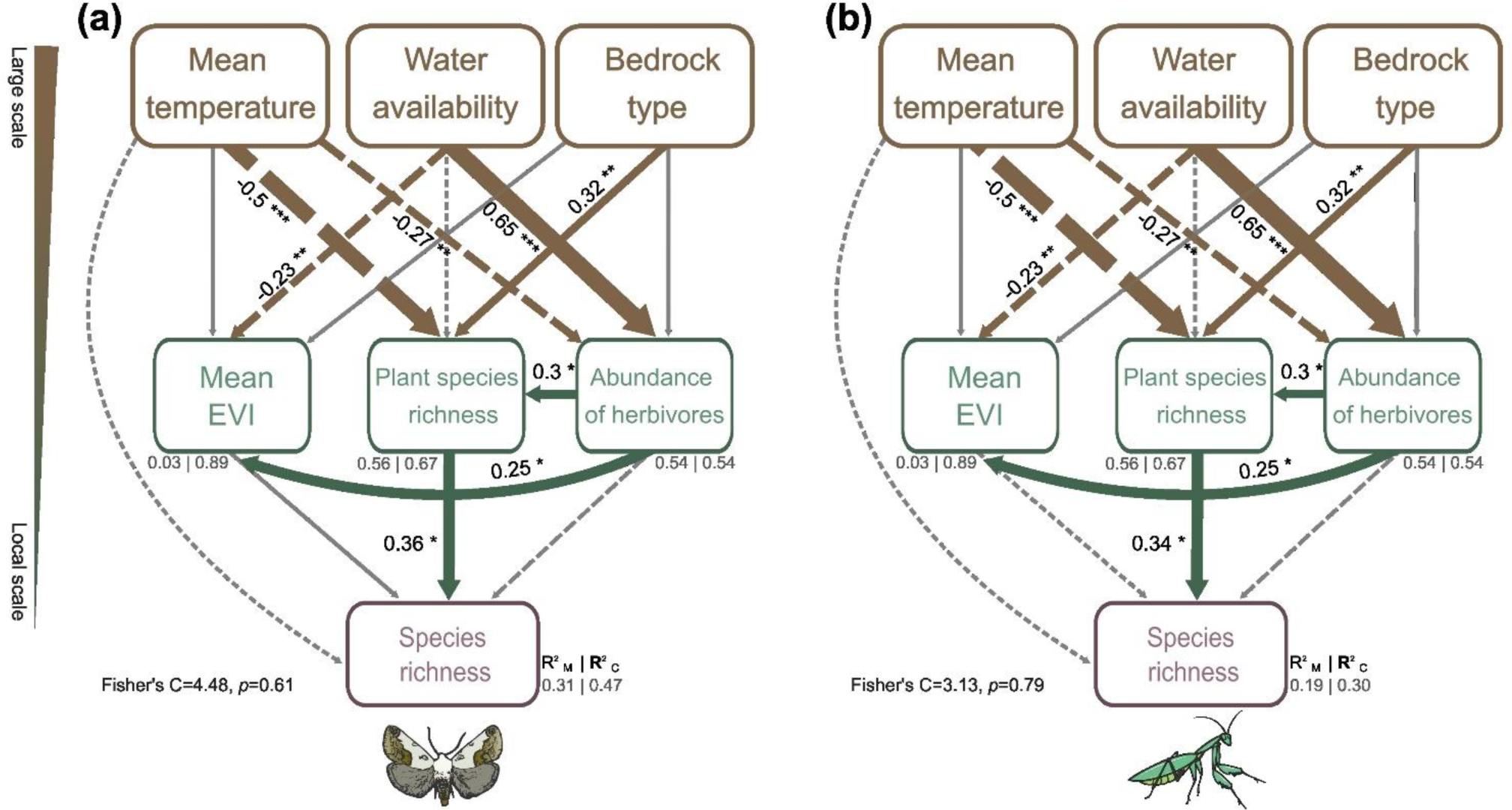
Path diagram illustrating how climate, habitat, disturbance, and vegetation variables affect species richness of (a) moths and (b) mantises in Kruger National Park, South Africa. Based on the results from piecewise SEMs, solid lines indicate positive associations, dashed lines indicate negative associations (*p ≤ 0.05, **p ≤ 0.01, ***p ≤ 0.001), and grey lines denote non-significant paths (p>0.05). Arrow widths are proportional to the standardized path coefficients, which are also noted alongside each arrow. Values in grey (R^2^) represent the proportions of variance explained by fixed (R^2^ marginal) and random effects (R^2^ conditional).

### Insect sampling

Both insect groups were sampled during the early (November) and late (February-March) wet seasons from 2018 to 2020. We employed portable light traps, each equipped with a dual-sided strip containing 48 ultraviolet (UV) light-emitting LEDs powered by 12V batteries (for details, see Delabye et al., 2022ab). These traps were operational throughout a complete night, from dusk to dawn, for each plot and season (i.e. each plot was sampled for a night in the early wet season and a night in the late wet season). To minimise the effects of weather and other potential unstudied temporal factors, the plots of each triplet were sampled during the same night. For more details on the sampling protocol, see Delabye et al. (2022b).

From the collected material, we identified all specimens of 12 moth families (Erebidae, Eutellidae, Noctuidae, Nolidae, Notodontidae, Eupterotidae, Lasiocampidae, Saturniidae, Sphingidae, Geometridae, Thyrididae, and Limacodidae), along with all mantis specimens. Both focal insect groups were identified based on the examination of external morphological characteristics and, when necessary, dissection of genitalia. Voucher moth specimens are stored at the Nature Education Centre of Jagiellonian University in Kraków, Poland, and at the Biology Centre, Czech Academy of Sciences, České Budějovice, Czechia. Voucher mantis specimens are stored at Department of Zoology, Faculty of Science, Charles University, Prague, Czechia.

### Environmental variables

The 60 sampling plots were characterized by 26 abiotic and biotic environmental variables divided into four groups (Table 1): *climate* (four characteristics of environmental temperature and two characteristics of rainfall), *habitat* (three water availability levels and two bedrock types), *disturbance* (two characteristics of fire history and large herbivore abundance), and *vegetation* (five measures of species richness, four characteristics of vegetation community composition, and six surrogates for plant biomass, such as vegetation cover and EVI). While most variables were represented by continuous values, water availability was categorised into three ordinal levels (perennial rivers, seasonal rivers, crests), and bedrock type into two categories (basalt and granite).

Vegetation data from the 60 studied plots were collected through botanical sampling during the rainy seasons (January to February in 2019 and 2020) when most plant species are identifiable (see Hejda et al., 2022 for more details). All vascular plant species in each plot were recorded, and their abundance was visually estimated using the Braun-Blanquet cover-abundance seven- grade scale; with the scores converted into percentage cover values for each species (Hejda et al., 2022). Vegetation composition was represented by the first four axes from a Principal Component Analysis (PCA) of the plant communities.

To estimate the abundance of large herbivores, camera traps (Bushnell Essential E3 Camera Trap with low glow IR flash) were installed in each plot, serviced approximately every three months starting in June 2018. Here we use animal records over 140 days, covering both the dry (August– October 2018) and rainy (December 2018–February 2019) seasons. We considered 18 species: buffalo, bushbuck, common duiker, elephant, giraffe, grysbok, hippo, impala, kudu, nyala, black rhino, white rhino, sable antelope, steenbok, tsessebe, waterbuck, wildebeest, and zebra (Hejda et al., 2022). The large herbivore abundance per plot was quantified by summing the records of all the herbivore species.

Given the strong correlations among some variables, we conducted correlation analyses to explore relationships between variables within each group. For variables that were highly correlated (r≥0.6, see Appendix S1: Figure S2), we retained those most meaningful for ecological interpretation of their potential relationship with the focal insect communities. To identify potential collinearity among the preselected variables, we calculated the variance inflation factor (VIF). This procedure yielded *mean temperature* as the only variable characterizing climate, *plant species richness* and *mean EVI* characterized vegetation, and the *mean number of fires* and *large herbivores abundance* characterized disturbance. Habitat remained represented by *water availability* and *bedrock type*.

### Drivers of insect species richness

To assess the direct and indirect relationships between the species richness of each insect group and environmental variables, we employed piecewise structural equations models (piecewise SEMs; Lefcheck, 2016; Figure 1a). Unlike classic path analyses, piecewise SEMs allow for independent solution of each model component, providing flexibility in specifying path relationships (Lefcheck, 2016). We log-transformed all variables except for temperature, which underwent a square root transformation. Subsequently, continuous variables were scaled (mean = 0, SD = 1). In the SEMs, each triplet was treated as a random effect, and the distance between triplets (based on GPS coordinates) was modelled as a continuous function.

For both moths and mantises, we constructed an initial piecewise SEM (“Saturated model”; see Appendix S1: Figure S3) based on available literature and hypothesized individual relationships between insect species richness and the non-correlated environmental variables (see Figure 1a, and hypotheses in Introduction). The model was further simplified by removing non-significant terms (p > 0.05) through backward elimination if their exclusion reduced Akaike’s information criterion (AIC). The fit of the reduced model was assessed by goodness-of-fit statistics, selecting the final model with the lowest AIC values (Grace, 2006). The Fisher’s C statistic was used to calculate the goodness of fit of the final model, and Shipley’s test of d-separation verified the absence of any missing paths in the model (Shipley, 2000).

For each path, we reported standardized path coefficients, estimating the expected change in the response variable as a function of the change in the explanatory variable in SD units.

Additionally, we reported marginal R^2^ (variance explained by fixed effects) and conditional R^2^ (variance explained by both fixed and random effects) values. For SEM-related analyses, we used the “coefs”, “lm”, and “lme” functions from the “piecewiseSEM”, “nlme”, and “lme4” packages, respectively (Lefcheck, 2016; de Boeck et al., 2011; Bates et al., 2015).

### Drivers of insect community composition

To assess the relationship between the community composition of each focal insect group and the four groups of environmental variables, we applied distance-based redundancy analysis (db- RDA) using Bray-Curtis distances. This method was selected based on results from a detrended correspondence analysis (DCA; Lepš & Šmilauer, 2003). For db-RDAs, we utilised the selected variables as previously described, with the exception of using the vegetation composition variables (represented by the first two axes’ scores from PCA; Table 1) instead of plant species richness. These variables effectively characterize how vegetation changes along environmental gradients and are deemed more suitable predictors of insect community composition (Schaffers et al., 2008).

To identify environmental variables significantly influencing the community composition of each insect group, we employed individual dissimilarity matrices as response data and used the environmental variables (after a forward selection procedure) as explanatory data (Lepš & Šmilauer, 2003). Prior to conducting db-RDAs (including the forward selection of environmental variables), all variables were transformed and scaled as in the SEM analyses. To reflect the spatial arrangement of the plots into triplets, a split-plot permutation scheme was implemented: individual triplets were treated as whole plots, with individual plots within these triplets treated as split plots.

To examine the contribution of each environmental variable, we applied hierarchical partition analysis (HP; Lai et al., 2022). Additionally, we employed variation partitioning analysis (VP) to evaluate relative contributions of individual variable groups (habitat, climate, vegetation, and disturbance) to community dissimilarity. This analysis decomposed the total variance in community dissimilarity into the pure effect of each group of variables, the variance shared between one or more groups, and the unexplained variance (Lai et al., 2022).

## Results

In the 60 study plots, we captured and identified 65,593 moth individuals representing 817 morphospecies, and 3,511 mantis individuals representing 38 morphospecies.

Our SEMs revealed that climate (mean temperature), habitat (bedrock type), disturbance (large herbivore abundance), and vegetation (plant species richness) collectively accounted for a significant proportion of the variation in species richness of moths (31%; Figure 2a) and in mantises (19%; Figure 2b). Notably, plant species richness emerged as the only variable with a highly significant and positive direct effect on species richness of both focal insect groups (Figure 2-3; Appendix S1 Table S1).

**Figure 3.**
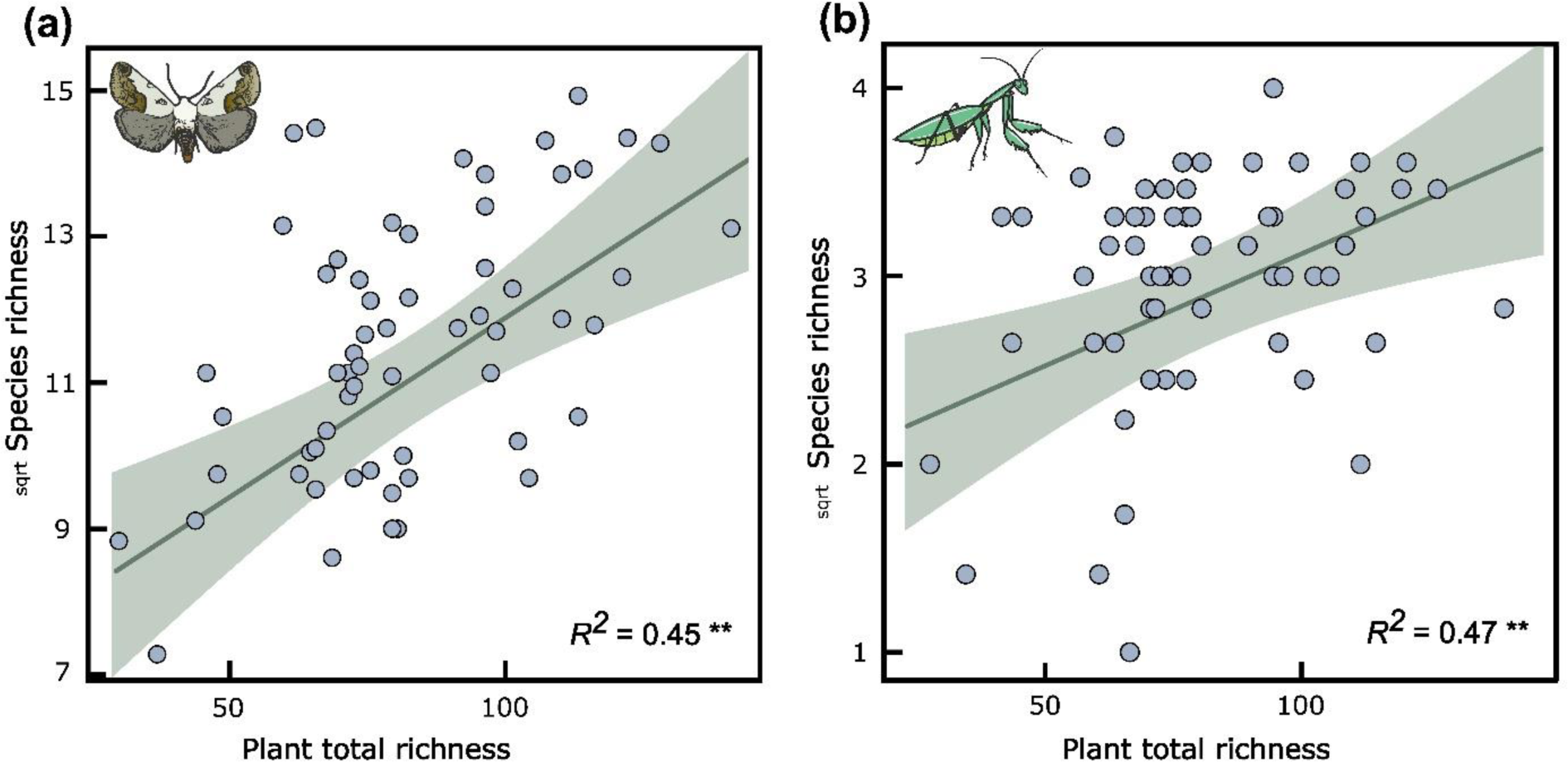
Effect of plant species richness on species richness of (a) moths and (b) mantises in Kruger National Park, South Africa. Lines and 95% confidence intervals represent predictions from linear mixed-effect models. ** *p* < 0.01

Additionally, climate, habitat, and disturbance characteristics had a significant direct effect on plant species richness in both final models (Figure 2ab). Specifically, mean temperature negatively influenced plant species richness and large herbivore abundance (Figure 2).

Moreover, bedrock type significantly influenced plant species richness, with basalt soils supporting a species-richer plant communities than granite soils. Furthermore, water availability demonstrated a significantly positive relationship with large herbivore abundance and mean EVI (Figure 2). Likewise, abundance of large herbivores positively influenced plant species richness and mean EVI (Figure 2).

According to the db-RDA models, all variables together significantly affected the species composition of moth (R^2^_adj_ = 16.2%) and mantis (R^2^_adj_ = 18.3%) communities. After forward selection, the final RDA models retained vegetation composition (PC1 and PC2) and water availability as predictors for both moth and mantis community composition. Additionally, mean temperature was retained for moths and bedrock type for mantises (Figure 4). The final models explained less variation in moth community composition (R^2^_adj_ = 8.3%) compared to mantis community composition (R^2^_adj_ = 10.6%).

**Figure 4.**
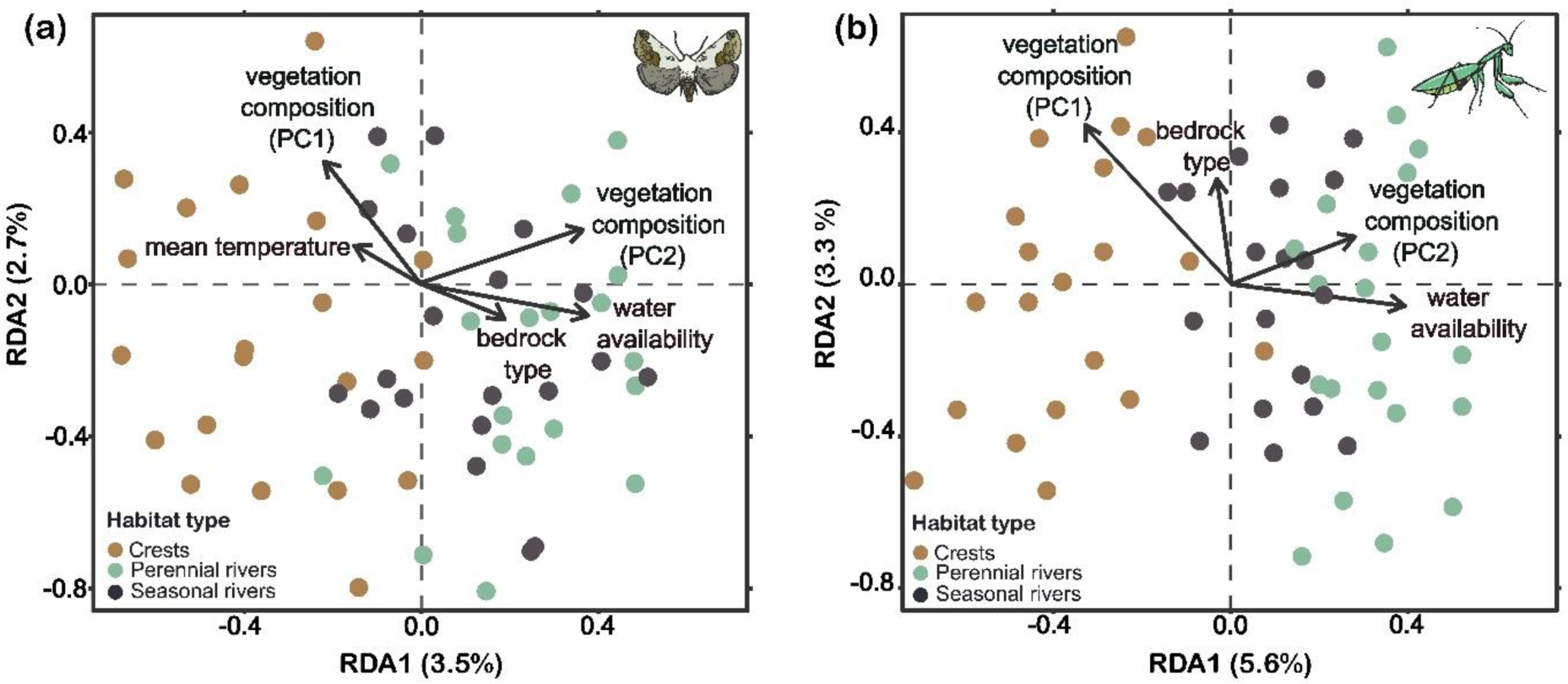
db-RDA ordination diagrams showing how species composition of (a) moth and (d) mantis communities relate to the forward-selected environmental variables.

The first axis of the db-RDA plot (RDA1) accounted for 3.5% of the total variation in moth and 5.6% in mantis community composition (Figure 4). RDA1 primarily delineated the differentiation between moth and mantis communities along the gradient of water availability.

Conversely, RDA2 contributed to 2.7% of the explained total variation in moths and 3.3% in mantises, which appeared to be related to changes in vegetation composition (Figure 4).

Hierarchical partitioning analyses indicated that among the vegetation variables, the composition (PC1) played a predominant role in community composition variation for both moths and mantises. The influence of mean temperature closely followed for moths (Figure 5ac), while all other variables accounted for less than 1% of the variation.

**Figure 5.**
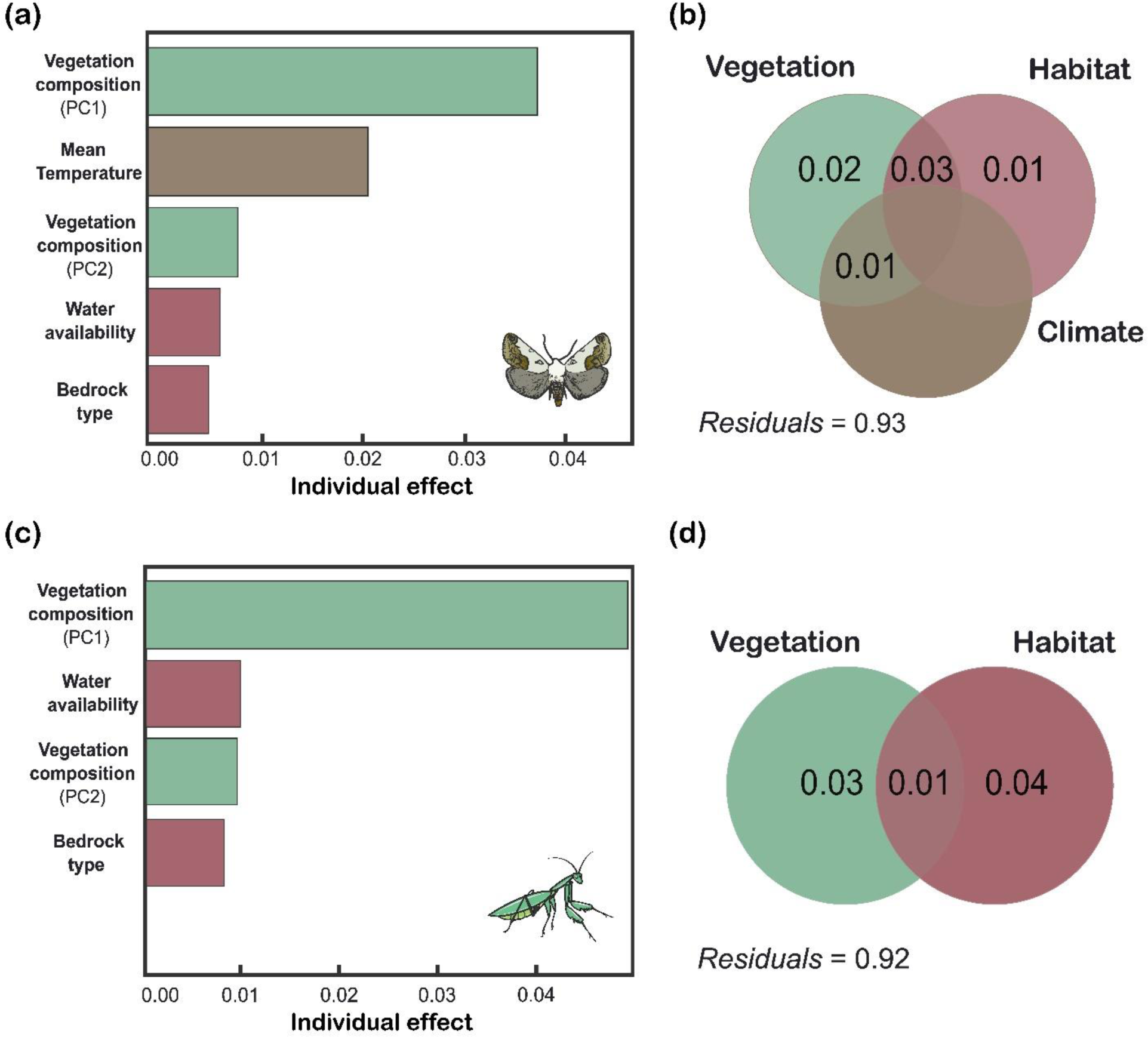
Hierarchical and variation partitioning of (a, b) moth and (c, d) mantis communities in terms of the percentage of variation explained by climate (brown), vegetation (pale green) and habitat variables (pale red). In the variation partitioning, the intersect represents the common contribution of the two group of variables, and blank spaces indicate 0 % of variance explained.

Variation partitioning analysis results indicated that the combined effects of vegetation and habitat variables accounted for roughly 1% of the variation in mantis and 3% in moth community composition (Figure 5bd). Additionally, the significant combined effect of climate and vegetation variables was only found for moths, explaining merely 1% of the total variation. Decomposing this variation revealed that vegetation variables alone accounted for 2% of the variance for moths and 3% for mantises. Individually, habitat variables were attributed to 1% and 4% of moth and mantis community composition, respectively (Figure 5bd).

## Discussion

Our study revealed complex impacts of climate, habitat, disturbance, and vegetation variables on insect species richness in the South African savannah ecosystems of Kruger National Park.

Notably, we confirmed a strong positive impact of plant species richness on diversity of phytophagous moth and carnivorous mantises, underlining the crucial role of vegetation in supporting diverse insect communities in savannah ecosystems. Furthermore, the effects of climate and habitat characteristics on insect diversity through plant species richness (Hejda et al., 2022) and large herbivore disturbance suggests complex interplays between environmental variables and dynamics of plant and insect communities, highlighting the nuanced nature of ecological interactions in these ecosystems (Jonsson et al., 2010; Leeuwis et al., 2018).

The hypothesised strong relationship between plant diversity and species richness of both insect groups is consistent with previous research of phytophagous insects (e.g. Kemp et al., 2017; Leone et al., 2023), as well as insect predators and parasitoids (e.g., Procheş & Cowling, 2006; Botha et al., 2016). This underscores that vegetation is crucial not only for phytophagous insect communities but also influences their natural enemies (Scherber et al., 2010; Balvanera et al., 2006). This also explains the virtually no differences in our results for moths and mantises. For phytophagous moth, our results can be related to the resource specialization hypothesis, predicting a correlation between diversity of consumers and their food resources (Keddy, 1984). The positive relationship between carnivorous insects and plant diversity may be related to the abovementioned increased diversity of herbivorous insect prey, but also to the higher complexity of species rich vegetation offering higher diversity of microhabitat conditions (Letourneau, 1987). Unfortunately, our study design does not allow separating these two mechanisms for any of the studied insect groups, especially as the influence of environmental variables on insect community composition was surprisingly low.

Our finding of no significant relationship between insect richness and proxies for plant biomass was unexpected. Previous studies of moths in South African savannahs (Delabye et al., 2022a) and various groups of herbivorous and other arthropods in KNP (Parker et al., 2023) suggested that primary productivity and vegetation complexity, both associated with plant biomass, are key drivers of insect diversity. However, these studies did not include plant diversity in their analyses of potential drivers. Based on our more robust dataset and detailed analyses, we propose that vegetation diversity may be a more important driver of insect diversity than plant biomass in South African savannahs.

Our results partially supported the hypothesis that climatic factors indirectly affect diversity of both insect groups through their impact on vegetation. While broader ecological theory and a part of empirical evidence emphasize the role of temperature in shaping insect diversity at a global scale (Mayr et al., 2020; Classen et al., 2015), our results aligned with the studies revealing the crucial role of plant communities in determining insect diversity at regional and local scales (Kemp & Ellis, 2017; Du et al., 2020). This pattern mirrors the broader dynamics observed in savannah ecosystems, where large-scale environmental factors influence local vegetation which shape insect communities (Bock et al., 2007; de Sassi et al., 2012; Zhu et al., 2014; Kemp et al., 2017; Shinohara & Yoshida, 2021). Although the negative effect of temperature on plant diversity through heat stress has been reported (Staver et al., 2017; Hejda et al., 2022), we rather expect the positive effect of precipitation which is strongly negatively correlated with temperature in KNP (Appendix S1: Figure S1). The crucial influence of precipitation on insect diversity in KNP has already been suggested by D’Souza et al. (2021). In our study with a broader spatial scope within the park, we revealed the mechanism of climate variables indirect impact on insect diversity through vegetation and disturbances.

The hypothesised indirect effects of habitat variables on insect diversity received varied support. Only bedrock type had a significant indirect effect on plant species richness, with basalt soils supporting higher plant diversity than granite (Hejda et al., 2022), which consequently enhanced insect diversity in our study. Water availability showed a different indirect mechanism of effects on insect communities. It influenced abundance of large herbivores which affected diversity of both focal insect groups through disturbing plant communities. Contrary to our expectations, a positive relationship emerged between large herbivore abundance and plant species richness, supporting previous studies that suggested mammal herbivore disturbances may facilitate species coexistence by creating diverse habitats and food resources for insects in savannah ecosystems (e.g., Bonnington et al., 2008; Wilkerson et al., 2013; Pryke et al., 2016).

### Conclusions and implication to conservation

Our findings emphasised the key role of vegetation in shaping insect diversity in the Afrotropical savannahs of Kruger National Park. They also highlighted the complex interactions among vegetation, climate, abiotic conditions, and disturbances from large herbivores, all of which have direct and indirect effects on insect diversity. Understanding these dynamics is crucial for devising effective conservation strategies, particularly in the face of climate change (Nooten et al., 2014). Our analyses revealed climate impacts insect communities predominantly through the negative correlations of temperature (and thus positive correlation of precipitation) with plant diversity. This underscores the vulnerability of savannah ecosystems to climate change, with significant implications for multiple trophic levels, encompassing both plant and insect communities.

Kruger National Park, along with the broader Southern Africa region, has been identified as highly vulnerable to climate change impacts (Engelbrecht et al., 2024; Rouault et al., 2024). Predictions indicate a decrease in precipitation during the rainy season, exacerbating the drying effect of elevated temperatures and leading to a pronounced decrease in soil moisture (Thavhana et al., 2024). These climate changes are expected to alter vegetation structure and composition, with consequences for insect communities. The decline in grass biomass reported during the 2016 drought in KNP already indicated the expected range of changing climate consequences (Wigley-Coetsee & Staver, 2020). Our results also emphasised the necessity of complex approach to conservation of African savannah biodiversity. Future studies should elucidate the mechanisms underlying the observed relationships by exploring finer-scale interactions and incorporating additional factors like plant-insect interactions and predator-prey dynamics (Hertzog et al., 2017). Integrating long-term monitoring and including variables such as water stress would significantly enhance our understanding of how temporal dynamics shape insect communities in response to changing environmental conditions.

Moreover, our study revealed the importance of seasonal rivers, with suitable compromise between water availability and disturbance intensity, among the fundamental conservation priorities. To protect insect communities, it is essential to support diverse vegetation through strategic management of large herbivores and regulated fires. Our study represents support of such practices successfully implemented in Kruger National Park over past decades (e.g., Robson & van Aarde, 2018; van Wilgen et al., 2021), demonstrating their effectiveness in maintaining insect biodiversity. Including seasonal rivers into these conservation practices seems to be an efficient strategy.

## Acknowledgements

We are grateful to Pavla Halamová and Markéta Staňková for help in the field; to Desmond Mabaso, Herman Ntimane, and Annoit Mashele for accompanying us in the field and keeping us safe; to Samantha Mabuza, Sharon Thompson, and Patricia Khoza for their assistance with arranging permits and other logistics in KNP. We used GPT-4 language model for English proofreading. This study was supported by the Czech Science Foundation (18-18495S and 21- 24186M).

